# Expanding P-NET, a multi-purpose biologically informed deep learning framework

**DOI:** 10.64898/2026.04.19.719454

**Authors:** Marc Glettig, Andrew Zhou, Chenzhang Zhou, Giuseppe Tarantino, Tyler Aprati, Eliezer M. Van Allen, David Liu, Haitham Elmarakeby

## Abstract

We present expanded P-NET, a versatile framework for deep learning in computational biology based on P-NET, leveraging biological pathways for interpretable predictions. Our framework achieves competitive performance in genomic & transcriptomic prediction tasks. We demonstrate its stability and interpretability compared to traditional machine learning models. P-NET 2.0 incorporates gene and pathway information, providing valuable insights into complex biological processes. The framework is publicly available, enabling its application to various computational biology tasks.

## 1. Introduction

In recent years, deep learning has emerged as a powerful tool for analyzing complex biological data and has revolutionized various domains of computational biology. The integration of deep learning techniques with biological knowledge has led to the development of biologically informed neural networks (BINN). These specialized neural network architectures are tailored to capture the intricate relationships within biological systems. One such architecture, the Pathway Neural Network (P-NET) (Elmarakeby et al., 2021), draws inspiration from the organizational principles of biological pathways and has shown promising results in biological applications (Hartman et al., 2023). Similar work has been done including biological and chemical knowledge to predict drug sensitivity (Deng et al., 2020).

In this work, we present an extended and versatile reimplementation of P-NET as a multi-purpose framework for deep learning applications in computational biology. The framework leverages the inherent structure and organization of biological pathways to limit the modeling capabilities of neural networks to meaningful connections. By incorporating domain-specific knowledge into the architecture, our framework is guided by the interpretability-by-design principle. Through this re-implementation and additional engineering, researchers can seamlessly apply deep learning techniques to a wide range of computational biology tasks, eliminating the need for task-specific network architectures. The flexibility and adaptability of our framework may not only save time and computational resources but also facilitate knowledge transfer across different biological applications.

To validate the effectiveness of our framework we first apply it to the original task of predicting metastatic vs. non-metastatic prostate cancer. Then we demonstrate the framework versatility and effectiveness by applying it to the clinically relevant task of predicting somatic mutations in cancer cell lines. Through extensive experiments on public datasets, we showcase the superior performance of our framework compared to regular deep learning architectures. Moreover, we provide an in-depth analysis of the learned representations and highlight the interpretability of our model, enabling biologists to gain valuable insights into complex biological processes.

## 2. Methods

### 2.1. Re-implementation of P-NET architecture

Our implementation of P-NET adopts a sparse feed-forward neural network architecture, Figure 1. The sparsity is achieved through masked linear layers, where the masks are generated based on the adjacency matrix of the Reactome hierarchical pathway database (Gillespie et al., 2022). The Reactome database provides a comprehensive collection of hierarchical pathway relations and gene-to-pathway associations.

**Figure 1.**
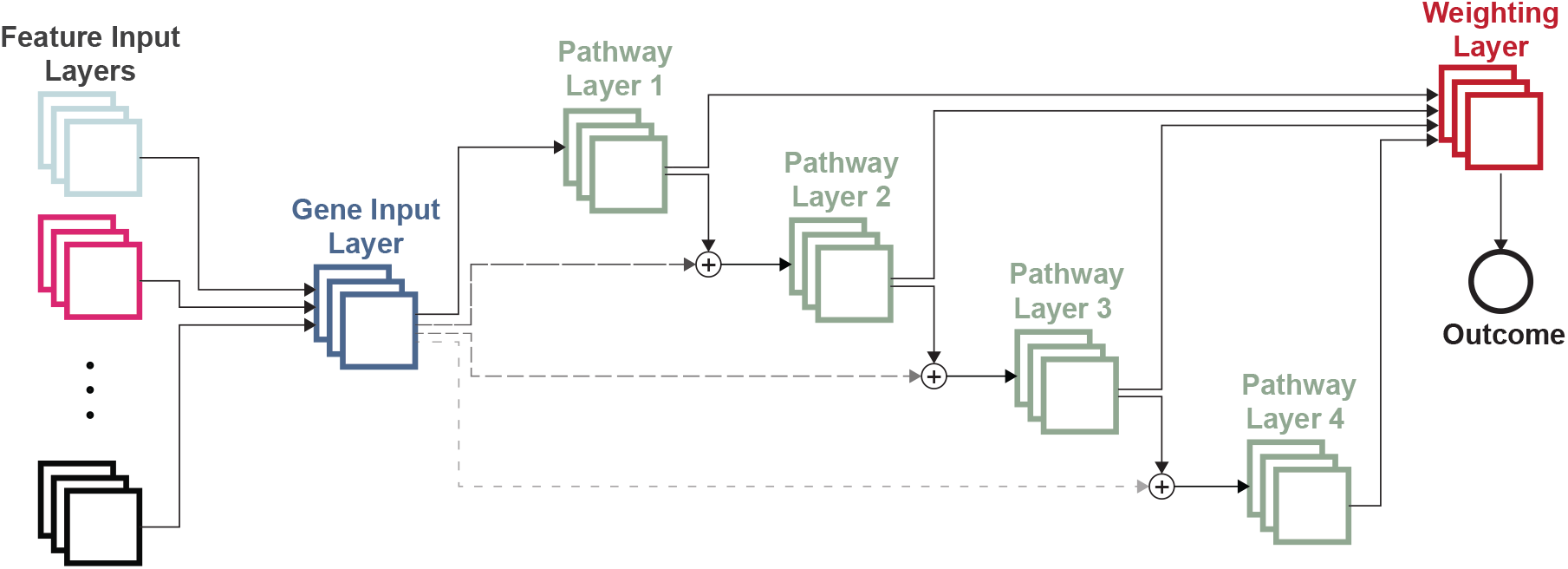
P-NET Model architecture; on the left we demonstrate how different gene level input modalities are connected to the Gene Input Layer (GIL). A single representation of each gene in the GIL is then connected to it’s respective biological pathways in the neural network layers. After each layer an outcome prediction is made and outcome predictions are then combined by a weighted average to form the final model prediction.

To handle different gene-level input modalities, we introduced a Gene Input Layer (GIL) that aggregates the input information for each gene. The GIL then establishes sparse connections between genes and their associated pathways.

The pathways, organized hierarchically, encompass various fine-grained biological sub-processes. By adding the pathway representations to the gene representations, we capture the collective influence of genes within higher-level pathways. It is important to note that each gene connects only to the lowest pathway level in which it is present.

Within each pathway layer, we incorporate a prediction head to generate predictions specific to that pathway layer. This property allows lower level pathways to directly and significantly impact the prediction. These pathway-level predictions are then combined using a weighted average approach, with the weights learned in a dedicated weighting layer. Finally, the overall outcome prediction is obtained as the weighted average of the predictions from different pathway layers.

### 2.2. Regulatory Layer

To further enhance the new P-NET model’s performance with RNA-sequencing data, we modified the architecture, by incorporating an additional “Regulatory Layer.” This layer, derived from the CollecTRI database, contains signed interactions between transcription factors and genes (Müller-Dott et al., 2023). The Regulatory Layer contains two types of connections: (1) a self-link for each transcription factor, and (2) a link from each gene to its regulating transcription factor, shown in Figure 2 and Figure 8. The output of the Regulatory Layer is summed element-wise with the output of the GIL and directed to the first Pathway Layer. This design ensures the preservation of gene-to-pathway interactions while augmenting them with gene-to-transcription factor interactions. Analogous to the Pathway Layers, the Regulatory Layer also contributes to the overall prediction through a dedicated prediction head. When fed RNA-sequencing data, the Regulatory Layer enables the network to effectively discern alterations in transcriptional regulation, potentially enhancing predictive accuracy and offering insights into aberrantly regulated pathways.

**Figure 2.**
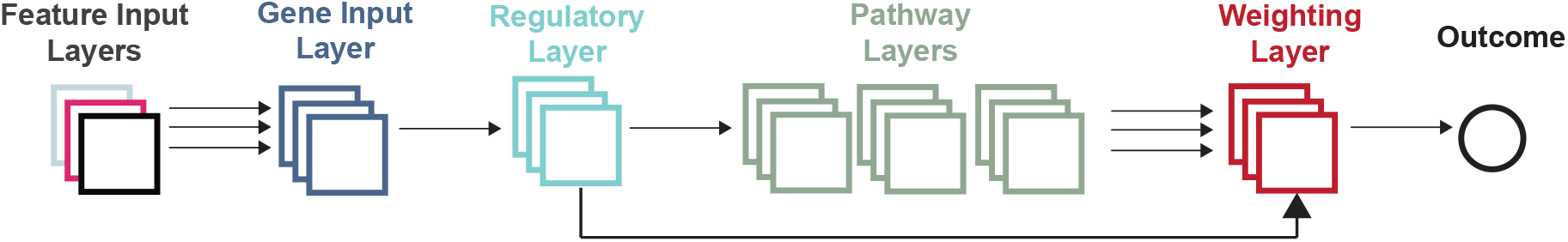
A variant of the P-NET Model with a Regulatory Layer. The Regulatory Layer contains connections from genes to the transcription factors that regulate them. This layer receives an integrated input from the GIL and outputs to the Pathway Layers and the Weighting Layer. Its output to the Pathway Layer is summed with the GIL output, and nodes are connected to Reactome Pathways in the same manner that GIL nodes are connected to the Pathway Layers in Figure 1.

### 2.3. Model performance assessments

To assess the performance of our model, we compared it against a normal feed-forward neural network, a sparse neural network with or without Biological knowledge, and traditional machine learning models such as random forests (RF) and support vector machines (SVM). The sparse neural network follows the same model architecture as P-NET, utilizing the same number of connections. However, the sparse neural network randomly selects the connections. This approach allows us to investigate the impact of structured connections derived from biological pathways versus randomly chosen connections in the model’s performance. An additional model we used for bench-marking is the Hallmark-Net, inspired by (Hao et al., 2018). Hallmark-Net uses the Hallmark gene sets (Liberzon et al., 2015) to define connections from the Gene Input Layer to the first sparse layer. Subsequent layers are identical to the sparse network. Compared to the random sparse network, Hallmark-Net bundles inputs from genes together into known gene-sets in the first hidden layer. Similar to P-NET this allows for biologically guided sparse connections in the model. These models were implemented and trained using established frameworks and libraries (Pedregosa et al., 2011).

We utilized the area under the receiver operating characteristic curve (AUC-ROC) to measure performance of the model. Additionally, to gain insights into feature and neuron importance scores, we employed the Integrated Gradients method (Sundararajan et al., 2017) and its extension (Shrikumar et al., 2018). These methods enabled us to attribute scores to individual features and neurons, highlighting their significance in the model’s decision-making process.

Furthermore, to assess the stability of feature importance scores, we introduced the concept of gene importance stability. Specifically, we measure the mean Jaccard index between top N gene sets across multiple runs. The Jaccard index is calculated pairwise, combining all possible pairs of runs. In A.5 we show how the number *N* of genes affect this score for multiple models in the metastatic prostate cancer prediction task. The best possible score of 1 is achieved if top N gene sets are identical across all runs. Given the results in Figure 10, we selected *N* = 20 to be the most sensitive in the prediction task at hand.

## 3. Results

### 3.1. Reproduction of Original Model

In this experiment, we aimed to validate the performance of our new P-NET framework by reproducing the outcomes from the original study (Elmarakeby et al., 2021). We compared the results obtained with our framework against various machine learning models. The evaluation was conducted on the original task of binary classification between metastatic and non-metastatic prostate cancer samples.

Our results demonstrated that all models performed reasonably well in capturing the metastatic vs. non-metastatic prediction task, with median AUCs across 10 folds exceeding 0.85. Specifically, our P-NET framework achieved higher AUC-ROC scores (median AUC = 0.91) compared to the other models, as shown in Figure 3. The original implementation of P-NET reported median AUC score of 0.93 across 5 fold-CV.

**Figure 3.**
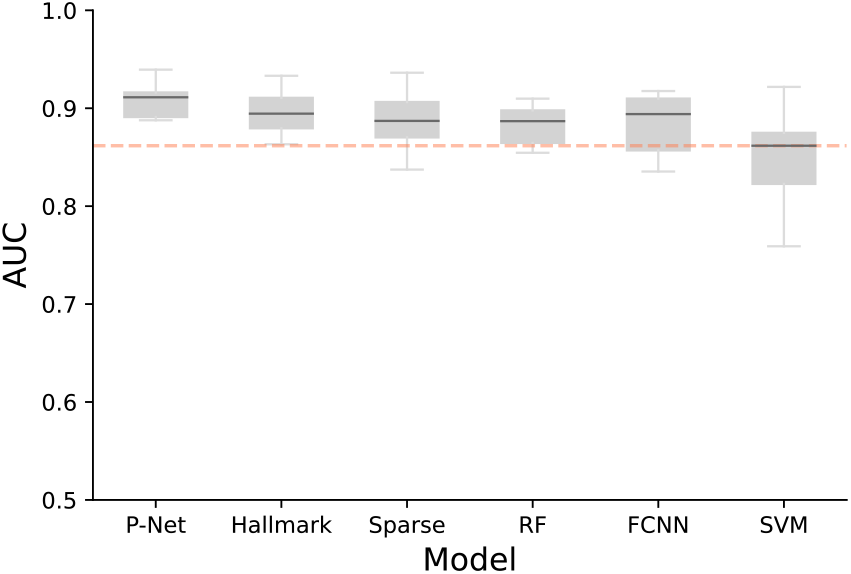
AUC (10-fold CV) for predicting metastatic vs. non-metastatic prostate cancer in the original cohort (Armenia et al., 2018). The median AUC for each model is represented by the black lines. Models are displayed in order of descending mean performance. Sparse deep learning models (P-NET, Hallmark and sparse NN) achieve similar performance to the traditional ML models (RF, FCNN and SVM).

Despite their lower performance, traditional machine learning models such as RF and SVM revealed high feature importance stability. We show how performance compares to feature stability in Figure 4. More complex models such as neural networks suffer from feature importance score instability. Sparse architectures like P-NET and the Random Sparse Neural Net can recover some of this instability, but still are far off from traditional models.

**Figure 4.**
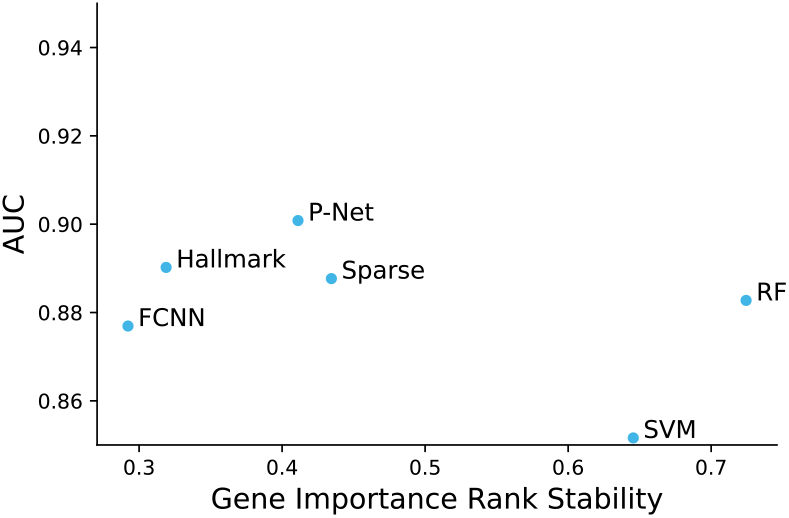
Comparison of variance in gene importance rank for the 20 most important genes. Rank stability is measured as the mean pairwise Jaccard index of the top gene set (see A.5). A score of 1 means perfectly stable importance values. RF & SVM, which capture simple interactions in the data, demonstrate robust interpretability. Among the deep learning models, sparse frameworks stand out as the more stable, with the highest stability scores.

In the analysis of node importance of hidden layers we show how P-NET sets itself apart from random sparse architectures. Traditional models like RF and SVM do not have such intermediate representations of the features. The interpretability analysis of the pathway layers revealed that our P-NET implementation exhibited more stable node importance scores compared to other deep models, such as the sparse neural network, as shown in Figure 5. This demonstrates that the embedding of biological information in the computational model may help enhance the predictions’ stability. Most importantly, the consistent importance scores for specific nodes elude to the fact the connections in P-NET represent relevant biological connections from Reactome. We performed further analysis of the signal to noise ratio (SNR), given by 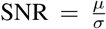, for the importance score of each pathway over the multiple runs. In Figure 6 the SNR of importance scores of the P-NET model exhibits a long tail compared to the random sparse neural network. This suggests that adding biological context and connections allows the model to identify important nodes within each layer that drive the phenotype. The mean importance score is lower for most hidden nodes in the P-NET architecture compared to the random sparse architecture. This can be attributed to the fact that P-NET focuses importance on meaningful pathways while importance is more evenly distributed in random architectures.

**Figure 5.**
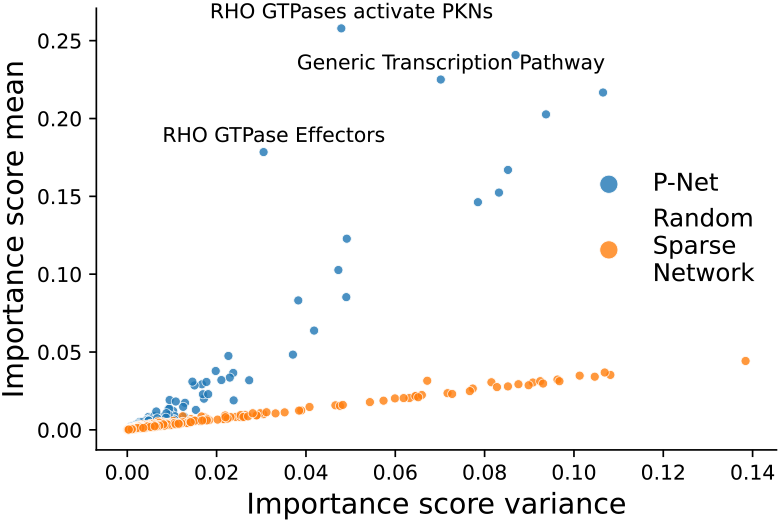
Scatterplot of all mean pathway importance scores vs. variance for P-NET and a random sparse neural net. Individual pathways in P-NET are able to consistently generate meaningful representations of the gene input date in the pathway layers, while pathway importance for the sparse network is random.

**Figure 6.**
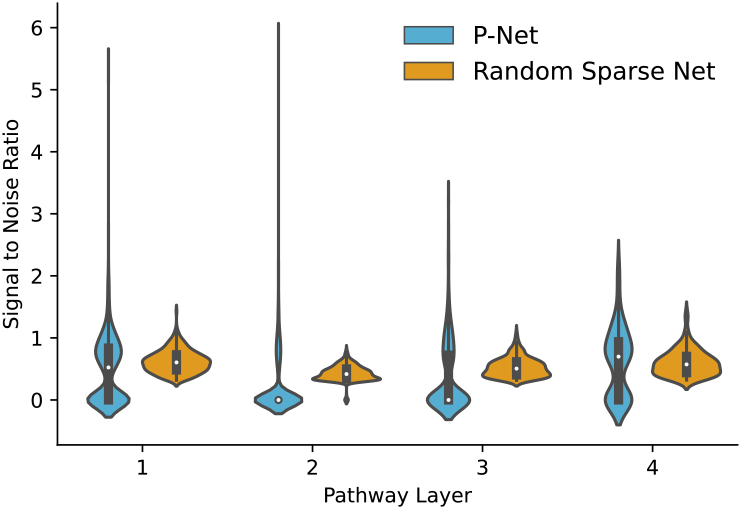
Violinplot of empirical signal to noise ratio 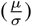 for pathway importance values across different layers by model architecture. Each violin plot also contains a interquartile boxplot (black) with the median value (white dot). We observe a similar distribution across the layers for the random sparse network. In P-NET, especially the lower level pathway layers contain multiple very consistently important nodes that likely correspond to meaningful biological entities.

### 3.2. Prediction of somatic mutations

In this experiment, we applied the P-NET framework to predict specific non-silent somatic mutations in cancer cell lines (CCLE) samples (Barretina et al., 2012). The selected mutations predicted are of high clinical interest and for some of them (TP53 & BRAF) the underlying biology has been exhaustively studied and described. This allows us to show that P-NET is able to recover previously described Biology.

By incorporating gene mutations and normalized RNA-seq data as inputs to the P-NET framework, we achieved accurate prediction of these mutations in CCLE samples. The performance of our model was evaluated using the AUC-ROC, which demonstrated competitive performance in distinguishing between samples with and without mutation of BRAF or TP53 (median test AUC of 0.96, 0.91 respectively). Additionally, we show that the regulatory layer allows to increase performance even more as well as making model performance more stable across runs as shown in Table 1 and Figure 7.

**Table 1.** Mutation prediction results in CCLE data. P-NET with regulatory layer outperforms baseline P-NET as well as a random forest model in the AUC-ROC score. Reported scores are median values across 10 fold CV for AUC as well as standard deviation across the ten folds to show performance stability.

| Mutation | Model | AUC | $\sigma$ |
| --- | --- | --- | --- |
| BRAF | P-NET | 0.96 | 0.04 |
|  | P-NET Reg | **0.97** | **0.03** |
|  | RF | 0.86 | 0.05 |
| TP53 | P-NET | 0.91 | 0.03 |
|  | P-NET Reg | 0.91 | 0.03 |
|  | RF | 0.87 | 0.03 |

**Figure 7.**
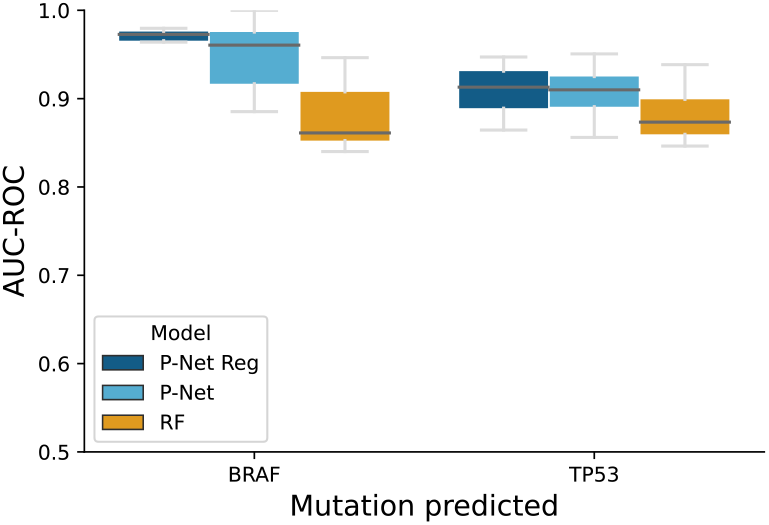
AUC (10-fold CV) for predicting non-silent mutations of genes of interest (BRAF, TP53) in CCLE samples. The training and testing were conducted on the TCGA cohort. P-NET and especially P-NET REG demonstrate strong predictive performance in discerning mutation events in CCLE samples compared to traditional machine learning methods. Furthermore, we show that incorporating the additional regulatory layer results in significant increases in performance stability.

**Figure 8.**
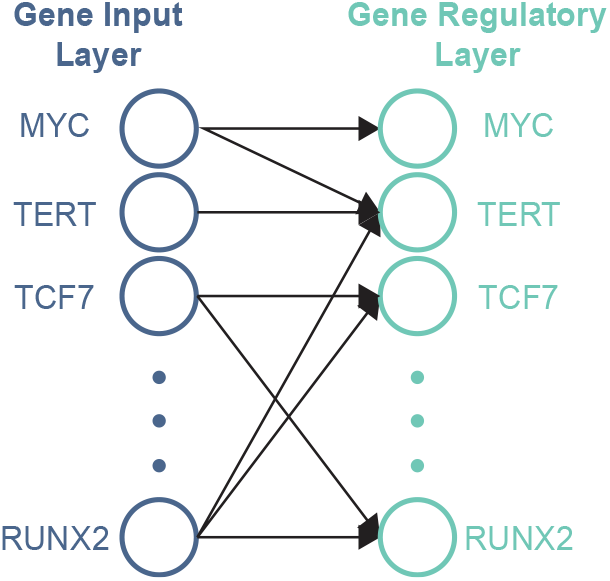
Detailed connections of the gene regulatory layer. Transcription factors connect to themselves as well as genes they have been described to regulate. The output of this regulatory representation is summed with the Gene Input layer before passed on to the pathway layers.

**Figure 9.**
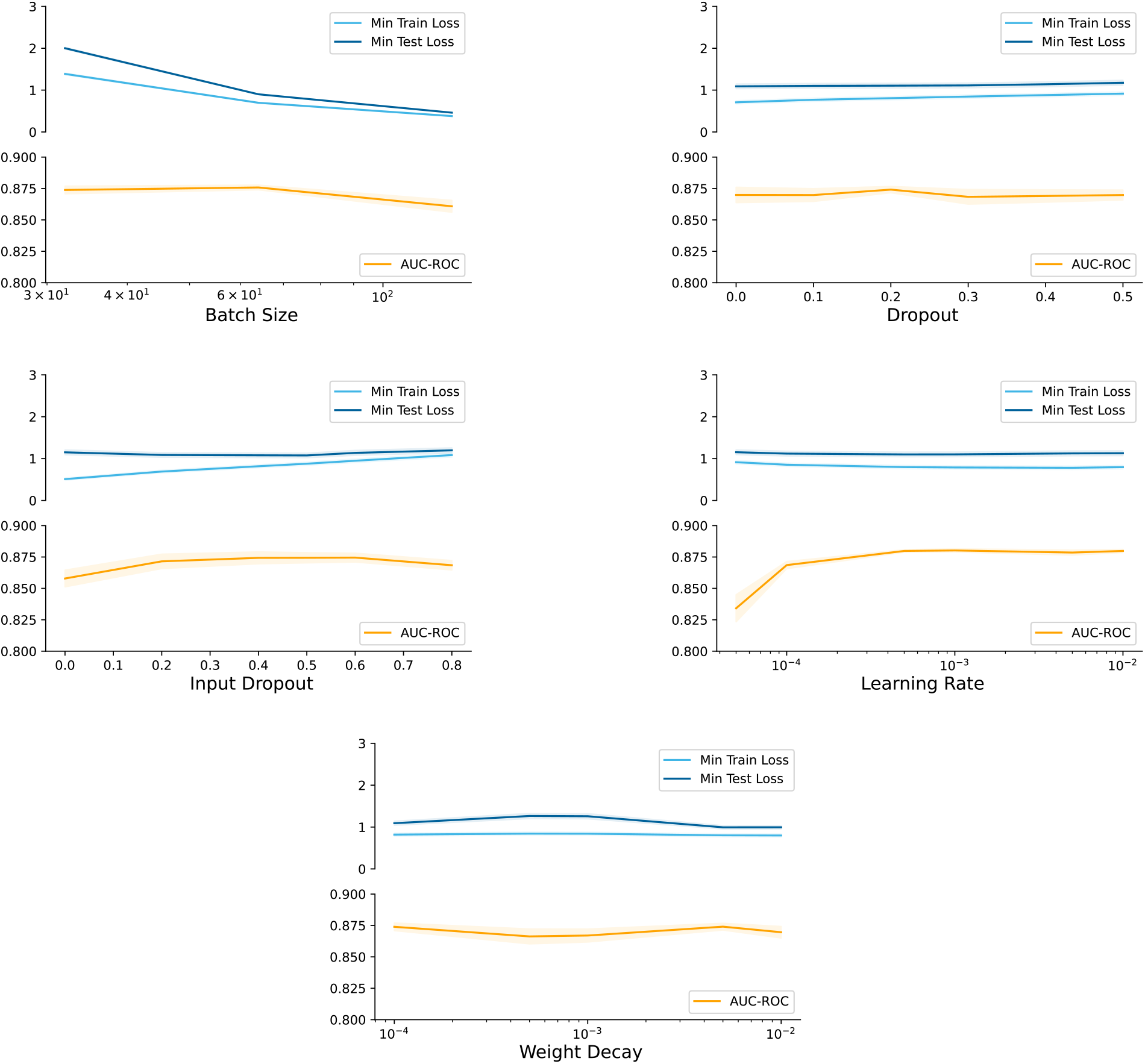
Hyper-parameter grid search has been performed with combinations of all different parameter. Here shown are mean results across axis displayed for batch size, dropout (in pathway layers), input drouput (in gene layers), learning rate and weight decay (L2-regularization). No significant performance decrease has been observed for any given hyperparameter other than a learning reate being too low. Hyper-parameter search has been performed on the task of metastatic vs primary prostate cancer.

The interpretability analysis allowed us to gain insights into the contributions of individual features and neurons in the prediction process. By leveraging the design of P-NET, it becomes feasible to link the importance scores to corresponding biological pathways and genes. Table 2 showcases the highest importance genes and pathways for the prediction of TP53 mutations. Notably, the most important gene used for prediction MDM2; is well described and an important negative regulator of TP53. Moreover, p21, encoded by the CDKN1A gene is a major target of p53.

**Table 2.** Top three significantly important Genes & Pathways per layer when predicting for TP53 mutation. Pathway/Gene significance are measured by the layerwise Z-score of their signal to noise ratio (SNR). We observe known p53 interacting genes as well as pathways describing these interactions being strongly significant.

|  | Layer | SNR | Z | p-val |
| --- | --- | --- | --- | --- |
| MDM2 | Gene | 5.88 | 10.20 | 7.83e-25 |
| CDKN1A | Gene | 4.36 | 6.98 | 1.50e-12 |
| BAX | Gene | 3.65 | 5.49 | 2.03e-08 |
| p53-Dependent G1 DNA Damage Response | Pathway 1 | 3.96 | 6.09 | 5.48e-10 |
| Release of apoptotic factors from the mitochondria | Pathway 1 | 2.96 | 4.30 | 8.68e-06 |
| Trafficking of AMPA receptors | Pathway 1 | 2.66 | 3.76 | 8.62e-05 |
| Transcriptional Regulation by TP53 | Pathway 2 | 3.93 | 7.12 | 5.57e-13 |
| p53-Dependent G1/S DNA damage checkpoint | Pathway 2 | 3.27 | 5.84 | 2.53e-09 |
| SUMO E3 ligases SUMOylate target proteins | Pathway 2 | 3.50 | 6.30 | 1.52e-10 |

Moreover, we note the abundance of pathways closely linked to TP53’s critical downstream functions within the pathway interpretability analysis. Among these, pathways crucial for cell cycle regulation, such as ‘p53-Dependent G1 DNA Damage Response’ and ‘p53-Dependent G1/S DNA damage checkpoint’, underscore the model’s sensitivity to biological processes involved in DNA damage repair and cell cycle inhibition. Additionally, the prominence of pathways like ‘Apoptotic factor-mediated response’ and ‘Transcriptional Regulation by TP53’ highlights the model’s recognition of pathways associated with TP53-induced apoptosis. These findings emphasize the model’s adeptness in capturing and leveraging biologically relevant pathways integral to predicting TP53 mutations.

## 4. Discussion

We successfully reproduced the outcomes from the original study using our new framework, which additionally provides more flexible entry points for new modalities inputs and architectures to encourage additional exploration and experimentation. The performance evaluation showcased competitive results, with our framework achieving performance superior to state-of-the-art models in identifying metastatic prostate cancer patients based on their genomic profiles. Notably, our P-NET implementation exhibited superior stability in hidden node importance scores compared to other deep models. This indicates that the incorporation of biological pathways in P-NET enables more reliable and interpretable predictions.

Furthermore, we applied the P-NET framework to predict oncogenes TP53 and BRAF in cancer cell line samples. Our choice to predict BRAF and TP53 mutations stemmed from their extensively documented roles as oncogenes in existing literature. These genes serve as well-established benchmarks due to the wealth of research surrounding them. By focusing on these well-studied genes, we aimed to assess whether our model’s identified important genes and pathways align with the established knowledge base. This approach allowed us to validate whether P-NET’s importance scores effectively coincide with widely recognized genetic markers, thus reinforcing the credibility of our model’s interpretability.

By integrating gene mutations and normalized RNA-seq data as inputs, our model accurately predicted oncogene mutations in CCLE samples. The results demonstrated superior performance in distinguishing between samples with and without mutation compared to RF, especially when using the additional regulatory layer. The interpretability analysis revealed a fascinating correlation: consistently important hidden nodes in our model directly correlated with biological processes intricately linked to the predicted mutated genes. Notably, this remarkable alignment underscores the model’s capacity to integrate Reactome network knowledge seamlessly into its neural architecture. The identification of consistently crucial hidden nodes associated with biological pathways directly connected to the predicted mutated genes stands as a testament to the model’s ability to intricately link its architecture with pertinent biological insights. This symbiotic relationship between the neural network’s internal representations and biologically relevant pathways demonstrates a profound integration of domain-specific knowledge into the model’s predictive framework.

The successful integration of domain-specific biological knowledge into the P-NET architecture not only elucidates insights into specific mutations but also signifies a promising methodology applicable across diverse problem domains. P-NET offers a robust framework to unearth novel and previously undiscovered biological relationships underlying various biological phenomena.

## 5. Conclusion

Our study showcases the versatility and effectiveness of the P-NET framework in deep learning applications within computational biology. By combining domain knowledge with advanced neural network techniques, our framework offers a unified solution to tackle various challenges in biological data analysis. Our framework provides a powerful tool for integrating multiple data modalities including genomic and transcriptomic profiles to predict biological and clinical outcomes. Moreover, our framework provides interpretable predictions and valuable insights into complex biological processes. This work contributes to the growing field of biologically-informed deep learning in computational biology, opening new avenues for leveraging biological knowledge to guide clinical prediction and biological discovery.

The P-NET comprehensive multi-purpose framework stands out for its user-friendly implementation and straightforward data integration. To facilitate its adoption, we have made the framework readily accessible at the following GitHub repository:

https://github.com/vanallenlab/pnet

Looking ahead, we aim to expand the scope of P-NET by exploring its applicability to unsupervised tasks and transfer-learning approaches in various cancer-related use cases. Additionally, we see potential in leveraging the framework for the analysis of single-cell data, further extending its utility and impact in computational biology.

## Acknowledgements

We would like to extend our sincere gratitude to the funding agencies, including the Department of Defense/LC220330, NIH K08CA234458, Doris Duke Clinical Scientist Development Award, Mark Foundation Emerging Leader Award, PCF-Movember Challenge Award, NIH U01CA233100, and NIH R01CA227388. Their generous financial support has been instrumental in enabling this research. We deeply appreciate their belief in our work and their investment in scientific endeavors.

Furthermore, we would like to express our appreciation to Dr. Jackson Nyman, Pasha Trukhanov, and Gwen Miller for their valuable contributions, insightful discussions, and constructive feedback throughout the development of this research project. Their expertise and collaboration have greatly enriched our work.

## A. Methods

### A.1. Dataset preparation

#### A.1.1. Metastatic Prostate Cancer data

For the validation of the original task, we followed the dataset preparation process described in (Elmarakeby et al., 2021). We utilized pre-processed data, which included mutation counts and copy number aberrations (CNA). The mutation data was transformed into a binary input format, while the CNA data was split into separate binary datasets representing amplification (AMP) and deletion (DEL). To ensure consistent evaluation, we employed the same data splits for 10-fold cross-validation, dividing the samples into training and test datasets. The target variable for this task involved binary classification between metastatic and non-metastatic prostate cancer samples.

#### A.1.2. Metastatic Prostate Cancer data

To showcase the application of our framework on an additional dataset and task we used the cancer cell line encyclopedia (CCLE). The data is publicly available under https://depmap.org/portal/download/all/. We used RNA-seq and mutation data from cell lines of all cancer types in the dataset. Mutations were only considered if non-silent and the respective gene predicted was removed as a feature from the input dataset.

### A.2. Regulatory layer

In the regulatory layer, genes from the input layer connect to all genes they regulate as given by the CollecTRI database. Most often genes would connect to themselves as well as other genes and regulatory embeddings would be added to the Gene Input Layer embedding of the feature space.

### A.3. Model training

The model was implemented using the PyTorch framework (Paszke et al., 2019). We initialized the weights using the He initialization method (He et al., 2015) and employed the Adam optimizer (Kingma & Ba, 2017) for training. A learning rate of 10^*-*4^ was utilized, and the training process spanned 300 epochs with the option of early stopping. Since all tasks involved binary prediction, Binary Cross Entropy was chosen as the loss function. To promote model generalization, we introduced L2 regularization with λ = 10^*-*3^.

### A.4. Hyper-parameter optimization

To optimize hyper-parameters as well as show robustness towards specific hyper-parameters, we performed grid search on batch size, dropout, gene dropout, learning rate and L2-regularization. We observe no significant drop or peak in performance for any given combination of hyper-parameters apart from too small learning rates as shown in Figure A.4. Default optimal hyper-parameters chosen and suggested for future users are a batch size of 64, dropout = 0.2, input dropout = 0.5, learning rate = 10^*-*4^ and λ = 10^*-*3^

### A.5. Gene stability

We assessed the stability of the gene importance for prediction by ranking the genes based on their importance score in each individual cross validation run. The generated 10 gene rank lists were then compared by using the Jaccard index of the top N gene set. The Jaccard index calculates the overlap of genes in two sets. We used the mean of the pairwise Jaccard index for all possible combination of runs to get a stability score for the gene importance. In figure 10 we show how the number of top genes considered *N* affects the stability score.

**Figure 10.**
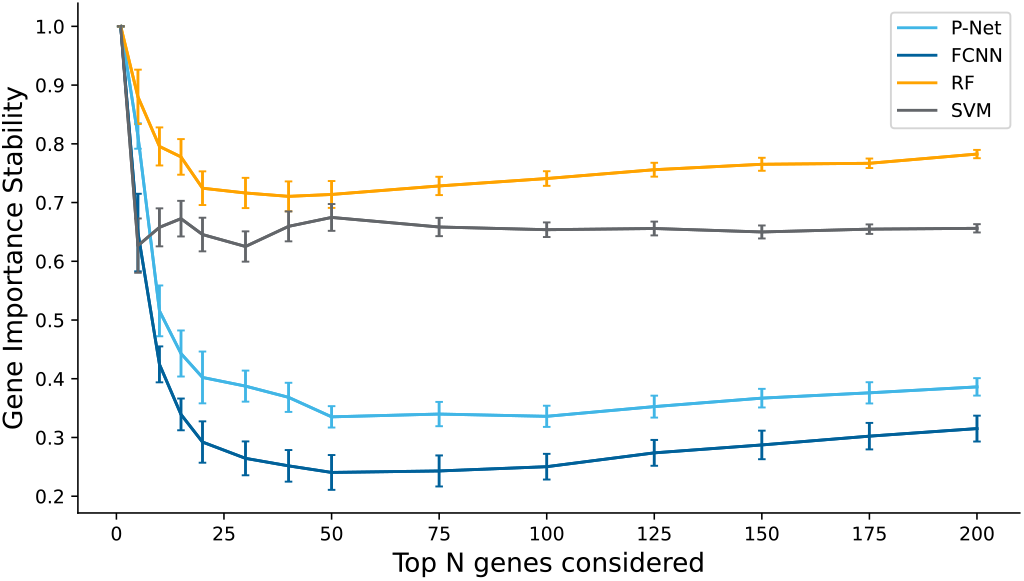
Mean pairwise Jaccard index over top N genes considered for stability measure. Scores start out at 1 for all models if only one gene is considered and drop to lower values as more genes are considered. Standard deviation measures obtained from bootstrapping.

### A.6. Mutation prediction

To validate our BRAF mutation prediction from cancer cell lines, we used the TCGA-SKCM cohort as external validation. Since the TCGA cohort contains sequencing from patient tissue, we expect a drop in predictive performance compared to the pure cancer line sequencing in CCLE. The model is still able to relatively well distinguish between samples with or without BRAF mutations as shown in figure 11.

**Figure 11.**
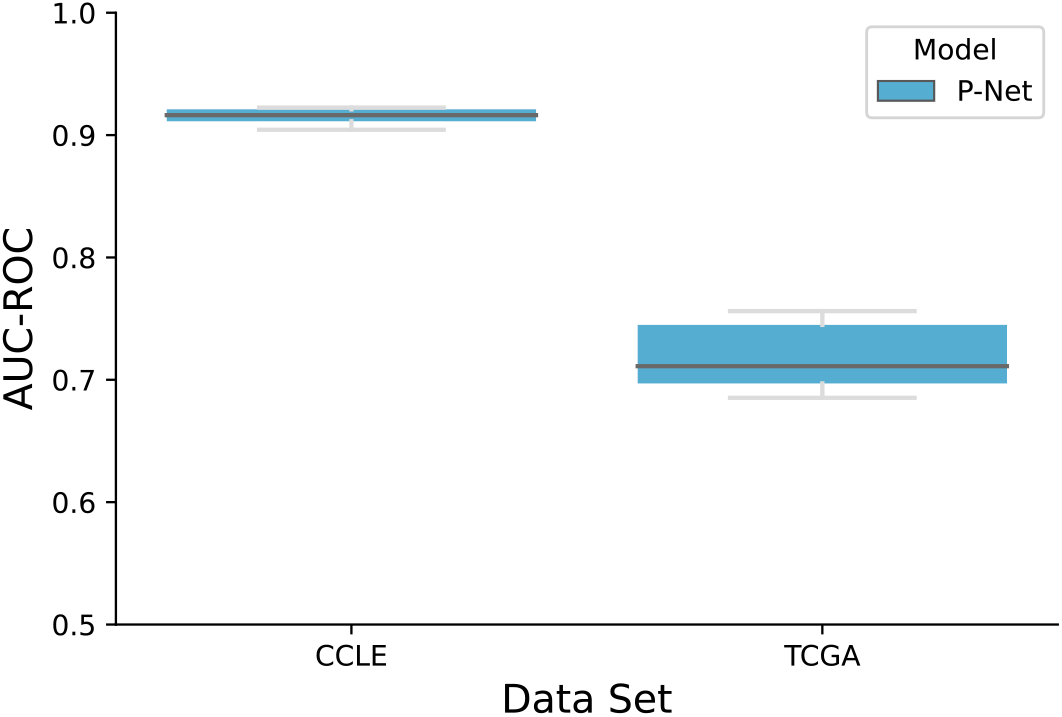
External validation of BRAF mutation prediction. Given the high prevalence of BRAF mutations in cutaneous melanoma, we used the TCGA-SKCM cohort as external validation cohort. Although we observe a strong performance drop, the model is still able to distinguish between mutated vs. non-mutated samples. Due to the TCGA being a clinical cohort rather than a cell line cohort this suggests our model identifies robust features to correclty predict BRAF mutations across model systems.

**Figure 12.**
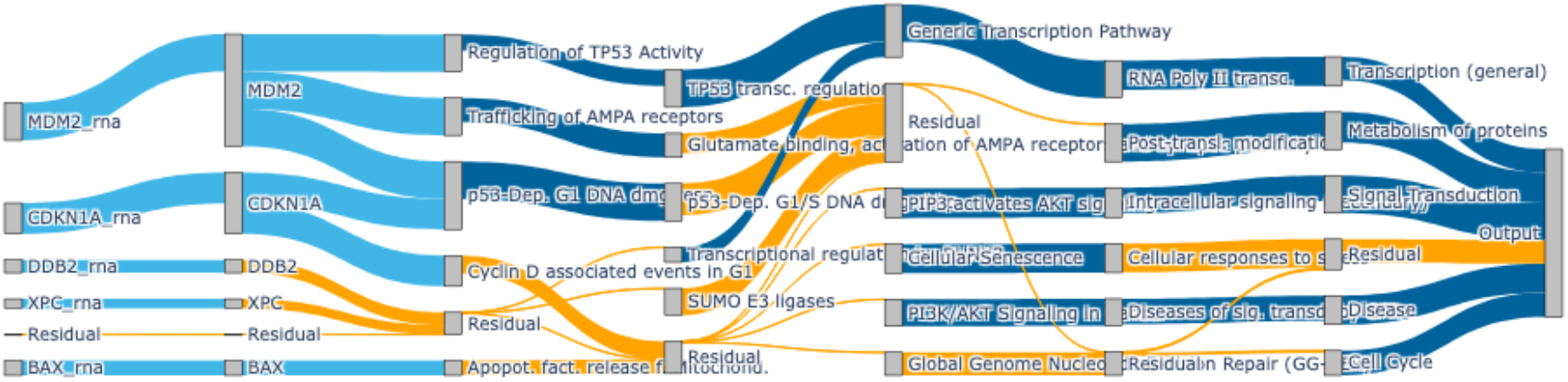
TODO: Update with less noisy sankey! Sankey diagram illustrating the flow of feature importances through P-NET in the prediction of nonsilent TP53 mutations. The diagram shows the most important genes and pathways per model layer. Notably, genes such as MDM2, CDKN1A and BAX, which have been heavily described in the TP53 pathway, as well several p53 driven pathways emerge as important drivers of the model’s prediction.

